# Local stabilization of subunit-subunit contacts causes global destabilization of Hepatitis B virus capsids

**DOI:** 10.1101/2020.01.24.918904

**Authors:** Christopher John Schlicksup, Patrick Laughlin, Steven Dunkelbarger, Joseph Che-Yen Wang, Adam Zlotnick

**Author notes:** Corresponding Author Adam Zlotnick, Molecular and Cellular Biology Department, Indiana University-Bloomington, Bloomington, IN 47401.

## Abstract

Development of antiviral molecules that bind virion is a strategy that remains in its infancy and the details of their mechanisms are poorly understood. Here we investigate the behavior of DBT1, a dibenzothiazapine, which specifically interacts with the capsid protein of Hepatitis B Virus (HBV). We found that DBT1 stabilizes protein-protein interaction, accelerates capsid assembly, and can induce formation of aberrant particles. Paradoxically, DBT1 can cause pre-formed capsids to dissociate. These activities may lead to (i) assembly of empty and defective capsids, inhibiting formation of new virus and (ii) disruption of mature viruses, which are metastable, to inhibit new infection. Using cryo-electron microscopy we observed that DBT1 led to asymmetric capsids where well-defined DBT1 density was bound at all inter-subunit contacts. These results suggest that DBT1 can support assembly by increasing buried surface area but induce disassembly of metastable capsids by favoring asymmetry to induce structural defects.

## Introduction

Hepatitis B Virus (HBV) remains a global health problem. In spite of an effective vaccine, more than 292 million people have chronic HBV, which can lead to cirrhosis, liver failure, and hepatocellular carcinoma and contributes to more than 880,000 deaths each year [1]. It is prevalent in Asia and has been described as a silent epidemic in Africa [2]. Roughly a third of people with chronic HBV will go on to develop severe liver disease [3]. However, at most 10% of people with chronic HBV have been diagnosed and less than 2% of them are being treated. Vaccine use drastically changes the demographics of HBV [4], but does not act on chronic infection. Nucleo(t/s)ide analogs, like Tenofovir and Entecavir, are effective at suppressing viral load and improve liver health, but do not cure chronic infection; once started, treatment may have to continue indefinitely due to the risk of viral flares [5]. Thus, there is a clear unmet medical need for new antiviral agents or treatment strategies which achieve a functional cure [6].

HBV is among the smallest human viruses. It is an enveloped virus with an icosahedral core containing the genome, a 3,200 base pair, relaxed circular dsDNA (rcDNA). The viral core protein (Cp) plays many roles in the viral lifecycle and is now being explored as a target for direct-acting antiviral agents. Once the virus loses its envelope during receptor mediated entry [7], the viral core protein becomes a major player. Cp mediates transport to nuclear pores by importin proteins [8, 9]. Cores bind to the nuclear basket via Cp and release viral DNA into the nucleus [10]. The viral rcDNA is “repaired” by host enzymes to yield the covalently closed circular form of the genome (cccDNA) [11, 12], which gains a cadre of nucleosomes and still has Cp bound to it [13, 14]. RNA transcripts from cccDNA are exported from the nucleus in spliced and unspliced forms [15]; a role for Cp in transcript export has also been proposed [16]. Once translated, Cp can assemble into either empty particles (about 90% of Cp [17, 18]) or RNA-filled cores that contain a pre-genomic RNA transcript (pgRNA) bound to viral reverse transcriptase [19]. The RNA-filled core becomes a metabolic compartment for synthesis of rcDNA from the linear pgRNA template. Of note, there are Cp mutations that affect synthesis of the second DNA strand, suggesting that the capsid plays an active allosteric role in this complicated reaction [20]. For the resulting DNA-filled capsids Cp can mediate transport to the nucleus, maintaining chronic infection, or alternatively Cp interacts with the viral envelope protein to initiate secretion from cells. These same fates are also available to empty capsids. Cp is conspicuously involved in most of the HBV lifecycle.

Many elements of Cp structure and assembly are well defined [21]. Cp has a 149-residue assembly domain and a 34-residue C-terminal nucleic acid-binding domain (CTD). The last nine residues of the assembly domain are sometimes referred to as a linker to the CTD [22, 23] and are largely disordered in structures. The assembly domain is dimeric with an intradimer interface formed by a large four helix bundle. The C-terminal fraction of the assembly domain, helix 5 and the subsequent loop and extended structure, form the interdimer contact in an assembled capsid. About 1200 Å^2^ of largely hydrophobic surface are buried at this interface [24]. Because the predominant form of HBV capsid has T=4 icosahedral symmetry (120 dimers), the interdimer interface must adopt four different, but quasi-equivalent, geometries. Despite the large buried surface, the interdimer contact energy is notably weak, about −3.1 kcal/mol at physiological ionic strength and stronger in higher salt [24]. Because subunits are each tetravalent, this weak energy corresponds to a pseudo critical concentration in the micromolar range. Since interaction energy is weak, errors during assembly can be thermodynamically edited out. Incomplete particles, overgrown particles, or aberrant assemblies can be trapped at higher ionic strengths, but can relax to the 120-dimer icosahedron over time and especially with lower salt concentrations as shown with single particle nanofluidics, single particle mass spectrometry, and SAXS [25-28].

The biophysics of assembly show how the reaction has adapted to work well in a limited range of conditions [29]. Therefore, pushing the reaction outside of that range using small molecules should be destructive to the virus. A Cp assembly-activating molecule may have three obvious activities [30, 31]. (i) A molecule that stimulates Cp to adopt an assembly-active state can lead to over-nucleation or premature nucleation, which for HBV could mean assembly without the reverse transcriptase-pgRNA complex. (ii) A molecule that stabilizes protein-protein interactions can also stabilize mistakes and lead to defective particles. (iii) A molecule that interferes with the normal protein-protein interaction geometry can lead to aberrant structures. These activities involve inducing structural changes and therefore should be stimulated by <u>c</u>ore <u>p</u>rotein <u>a</u>llosteric <u>m</u>odulators, or CpAMs. A CpAM that modulates assembly may also alter Cp’s ability to perform other activities.

Multiple CpAM chemotypes have been identified. Phenylpropenamides (PPAs) and heteroaryldihydropyrimidines (HAPs) were discovered in naïve screens and later shown to act via the Cp [32-35]. In both cases, these molecules are assembly agonists that accelerate kinetics and stabilize Cp-Cp interactions, though PPAs always lead to normal capsids while HAPs can create strikingly aberrant Cp polymers [36, 37]. Sulfamoyl benzamides (SBAs) show similar properties to PPAs, forming normal capsids [38]. A likely mechanism of action for these CpAMs is that they stimulate assembly leading to empty and defective particles (see [32]). Recently, an antifungal agent, ciclopirox, was reported to interfere with HBV capsid assembly in cells, and drive aberrant Cp polymer in vitro [39]. Each of these CpAM chemotypes bind at a similar location at the subunit interfaces but can produce very distinct outcomes in the products seen in assembly reactions. Even within a single chemotype, the products that form vary depending on substituents, stoichiometry, and solution conditions [40].

The differences in assembly product morphology have led some to classify CpAMs as either Type 1 molecules that drive aberrant assembly, or Type 2 molecules that don’t. In this work, we seek a more nuanced view of CpAM mechanism, where molecules can have multiple different activities, each of which are further dependent on the local environment. We present a novel chemical scaffold, dibenzothiazapine (DBT), which can paradoxically induce HBV capsid stabilization and destabilization. Assembly reactions can lead to both normal and aberrant capsids, and the molecule has a quasi-equivalent binding preference that is distinct from what is seen in other CpAMs. The effects of DBT1 on the physical chemistry of assembly provide a basis for interpreting and predicting the biological effects of CpAMs.

## Results and Discussion

### DBT locally stabilizes per-contact subunit interfaces

The HBV core protein has many functions, but its critical role is to assemble capsid and provide a stable compartment for reverse transcription. Eventually, the capsids must undergo disassembly to release the genome and infect a host cell. Interfering with assembly is the best characterized activity of CpAMs, which include the DBT chemotype. Though several hundred DBT variants have been identified [41], their physical chemistry has not been well characterized. Here we examine the behavior of DBT1 (**Fig. 1a inset**). First, we ask whether DBT1 modifies assembly kinetics. In vivo, Cp assembly is triggered by a genome-polymerase complex to eventually to yield virus [42], or spontaneously to yield empty capsid [17]. CpAMs can stimulate assembly to increase the yield of empty and aberrant capsids [37]. Assembly kinetics were monitored using 90° light scattering, where the scattering signal is proportional to the size and number of assembled products (**Fig. 1a**). In solution conditions where capsid assembly is normally undetectable, the DBT molecule promotes assembly in a robust and dose dependent manner. Once the reaction approaches a steady state equilibrium, the fraction of assembled product can be estimated from size exclusion chromatography (**Fig. 1b**). This comparison addresses the question of whether DBT1 affects thermodynamics as well as kinetics. Here we see that DBT promotes assembly in a dose dependent manner, shifting the reaction equilibrium away from free subunit and towards formation of larger structures. In the context of an assembly reaction model, these observations are consistent with a strengthening of the per-contact energy, ΔG_cont_, of subunit-subunit association. This may be accomplished by inducing Cp to enter an assembly-active transition state and/or by stabilizing Cp-Cp interactions. To quantify the effect of DBT1, we use the ratio of capsid to dimer, an equilibrium constant, to determine the change in ΔG_contact_ as a function of [DBT1]. From these data we observe that DBT1 strengthens association energy from −3.1 kcal/mol in 150mM NaCl to asymptotically approach −4.4 kcal/mol **(Fig. 1b inset).** The apparent equilibrium constant for induced assembly is 0.59µM, indicating that DBT1 reversibly binds capsid during assembly.

**Figure 1.**
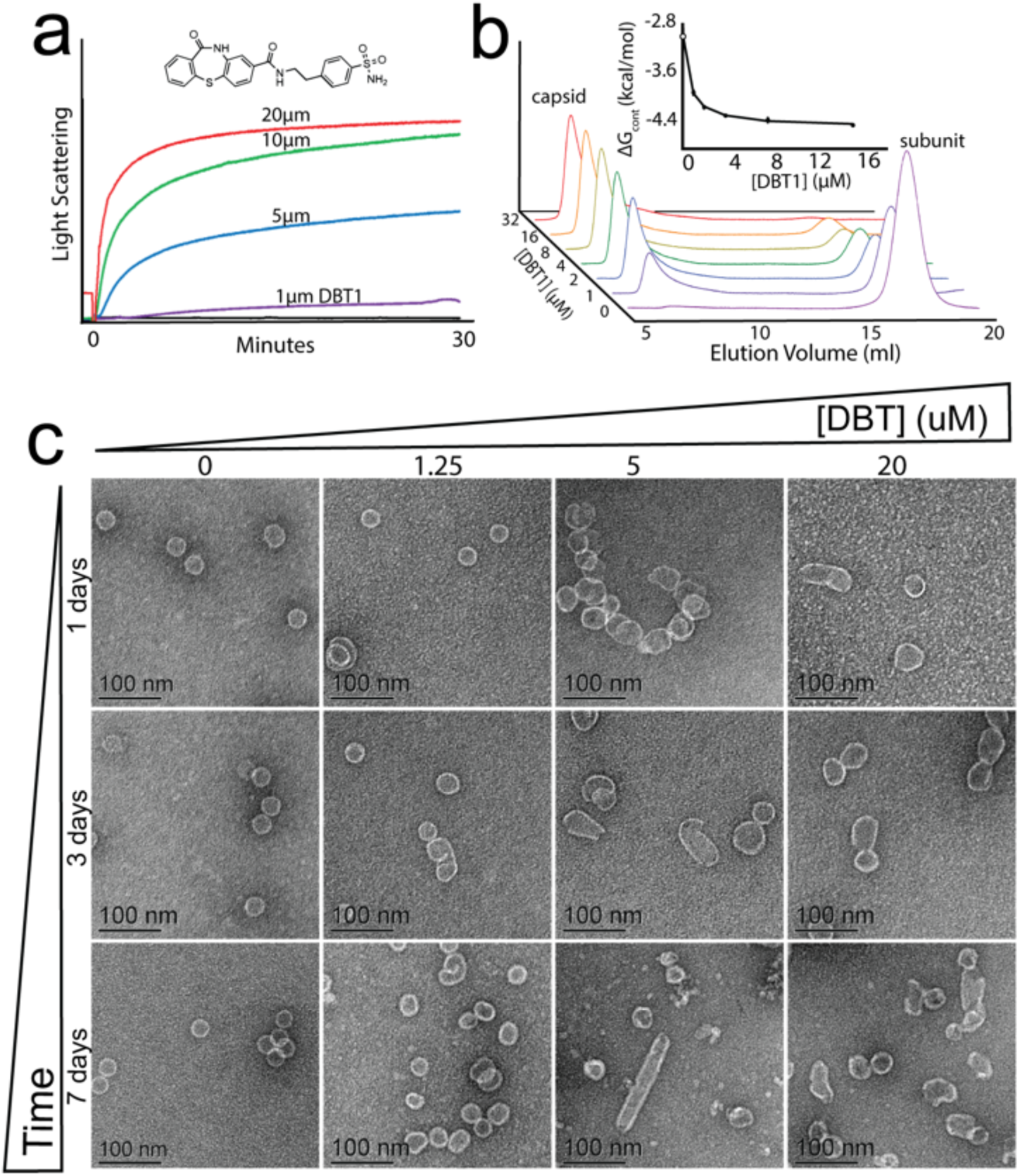
DBT accelerates assembly and activates mis-assembly. **(a)** Reactions of 5µM Cp149 assembled in 150 mM NaCl (final concentrations) are monitored by 90° light scattering. The increase in scattering signal that appear with increasing DBT1 concentration (structure inset) indicates the formation of capsid and aberrant structures. In the absence of DBT1 (black line), assembly is undetectable at these conditions. **(b)** Size exclusion chromatography (SEC) confirms assembly and quantifies assembled products and free Cp149 dimer. Both methods indicate that DBT1 increases assembly relative to the control (absence of DBT1, black). The inset in panel **(b)** fits the association energy per subunit contact (ΔG_cont_) versus [DBT1] to a hyperbolic binding curve. The reference value for Cp149 without DBT1 (white circle) was previously determined [24]. **(c)** Reaction products visualized by TEM correlate with light scattering. Over time, assembly products rearrange to form increasingly aberrant structures.

Because the amount of light scattering at high DBT1 is more than might be expected from a drug free reaction, and because the stronger association energy would decrease the potential for thermodynamic editing [29], we examined reaction products for defects by electron microscopy (EM). We observed a dose-dependent appearance of larger non-capsid structures (**Fig. 1c**). Notably, even at sub-stoichiometric DBT1 concentrations (1.25µM), some particles were distinctly aberrant. This suggests that a defect in the growth of those particles occurred early during assembly, perhaps at nucleation. At high DBT1 concentrations, a larger number of progressively larger misshapen particles appeared; nonetheless, many particles with wild type morphology appear simultaneously. Concentration-dependent mis-assembly is also characteristic of HAPs, where sub-stoichiometric concentrations lead to normal capsids and higher concentrations are required to produce larger, sheet-like complexes [43]. A distinctive activity of DBT1 is the slow rearrangement of reaction products with time. At the scale of several days, the fraction of icosahedral particles decreases, while the fraction of large poly-disperse structures increases. This maturation effect is perhaps most striking with the appearance of cylindrical structures which appear at high DBT1 concentrations and long times. This is significant because it suggests that the initial assembly products are not actually at thermodynamic minima, but rather are kinetic traps. By observing the behavior of these reactions, we gain an understanding of what might be happening during particle formation and maturation. At early times, more particles are icosahedral; at longer times particles become more aberrant. This indicates that the kinetic barrier to form spherical particles is relatively low. We note that in vivo, RNA-filled particles are very stable whereas mature particles filled with relaxed circular dsDNA are predicted and observed to be unstable [44, 45]. Paradoxically, we have shown that DBT1 stabilizes Cp-Cp interactions while destabilizing icosahedra.

### DBT1 globally destabilizes capsids

We hypothesize that rearrangement of smaller, spherical structures into larger, preferentially cylindrical structures occurs by transiently releasing subunits. To test for the appearance of free subunit, we added CpAMs to a homogenous population of pre-assembled capsids and examined the complexes by native agarose gel electrophoresis (**Fig. 2**). Native agarose gel electrophoresis (**Fig. 2a**) can resolve subtle differences in the size of assembly products (>3-4MDa) and also resolve free dimer (35kDa). Electrophoresis also places capsids into conditions where they are metastable and could never spontaneously assemble to favor transient dissociation. By separating dimer and capsid (or other larger oligomer), we also prevent reassociation, allowing us to visualize the transiently released subunits.

**Figure 2.**
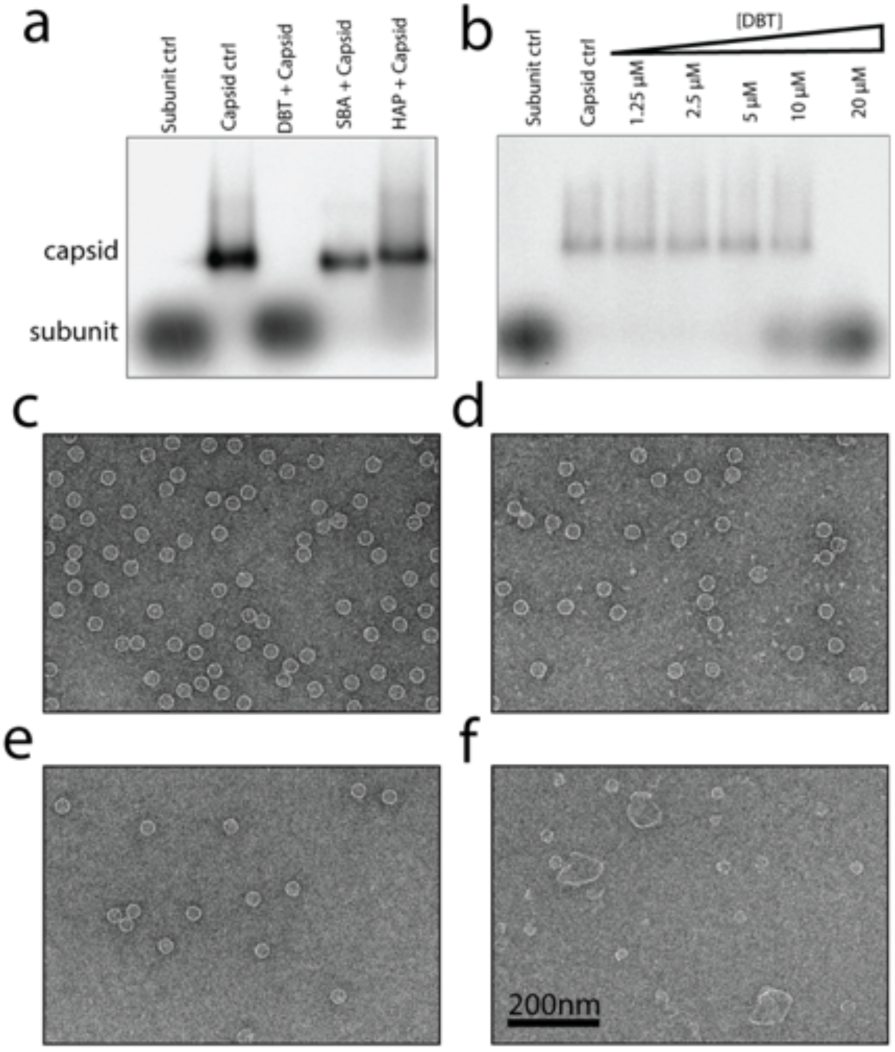
DBT induces capsid destabilization. **(a)** Native agarose gel electrophoresis can resolve both intact capsids and free subunit. We use an assembly incompetent mutant, Cp149-Y132A as a control for free dimer. Preformed capsids treated with a 20µM DBT1 (4:1 molar ratio of DBT to Cp) are quantitatively destabilized and co-migrate with free subunit. In contrast, two other capsid-directed molecules (SBA, HAP) show very different behaviors. SBA-treated capsid migrates with untreated capsid. The band for HAP-treated capsids smear broadly to indicate the presence of products that are capsid size or larger. **(b)** The transition from capsid to subunit is concentration dependent, with the capsids fully dissociated only at a molar excess of DBT1 (here 5 µM subunit results in 10 µM of subunit interfaces). **(c-f)** The reactions shown on the gel in panel **(a)** are visualized by TEM in panels. **(c)** Untreated capsids provide a reference point. **(d)** In a non-perturbing environment, most capsids treated with DBT appear normal. Some capsids displayed what may be broken regions or holes. **(e)** SBA treated capsids look identical to untreated capsids. **(f)** HAP treated capsids become disrupted, forming larger structures., including many particles which are capsid sized, but appear distorted. The presence of the larger particles suggests a process of disruption followed by reassembly..

We chose three molecules (DBT1, SBA0013, and HAP12) to represent three of the CpAM chemotypes (DBT, SBA, and HAP), each of which exhibits distinct behaviors during assembly reactions. Introducing DBT to pre-formed capsid causes all protein to co-migrate with free subunit, indicating that under these conditions the molecule induces complete disassembly. In a titration of capsid by DBT1 we found that quantitative dissociation was observed at 20µM DBT1 (with 5µM capsid) while at 10µM DBT1 little dissociation was observed. From the assembly-based titration (**Fig. 1b**), this suggests that we need almost every site filled (94% filled at 10µM versus 97% filled at 20µM, assuming the simple binding model of Fig. 1b, inset) to drive dissociation. Upon inspection by EM (**Fig. 2d**), we found that under equilibrium conditions the protein remains largely in capsid form. However, reactions eluting slowly over size exclusion chromatography recapitulate the dose-dependent disassembly seen by gel electrophoresis **(Fig. S1b).** This suggests that dimers were transiently released and that the resulting capsids with defects then unraveled during elution.

In comparison, the SBA-treated capsids co-migrate with apo-capsid and look indistinguishable from the control by electron microscopy (**Fig. 2e**). SBAs also drive assembly and yield morphologically normal capsids; the persistence of capsids under the equilibrium-perturbing conditions of electrophoretic separation indicate that few dimers are released and suggests that drug-bound capsid is the lowest energy state. Capsids treated by HAPs also change their migration, leading to smearing of the capsid band to progressively slower migrating species. This suggests that Cp dimers are released and reassemble into large structures reminiscent of what would be expected for HAP-directed mis-assembly starting from free subunit (**Fig. 2f, S1c**). The extent of smearing by the HAP was sensitive to the time and concentration of HAP used. We find it of great interest that DBT1 is sufficient to completely disrupt capsids during electrophoresis but insufficient to rearrange them. In contrast, the HAP appears strong enough to keep new assemblies together on the gel, but it should be kept in mind that disruption and re-assembly occurred before electrophoresis. In effect, DBT1 has poised the capsids for disassembly.

### Cryo-EM Structure

To gain insight into the mechanism of the drug-induced stabilization and destabilization, we carried out cryo-EM reconstructions of pre-formed Cp150 capsids treated with DBT. The Cp150 construct incorporates a C-terminal cysteine that allows Cp dimers to crosslink around fivefolds and quasi-sixfolds to prevent capsid dissociation or rearrangement [46]. This crosslink also minimizes drug-induced excursions from icosahedral symmetry. Initially we imposed icosahedral symmetry during image reconstruction (**Fig. 3a**) as historically this has been an accurate assumption and enhances signal-to-noise ratio by 60-fold averaging. Quasi-equivalence explains how chemically identical subunits fit into structurally distinct, but similar, locations in an icosahedrally symmetric structure. For HBV, with T=4 quasi-equivalence, the 240 interfaces have four unique conformations, labelled A, B, C, or D. In our icosahedral reconstruction we saw density that could be attributed to DBT1 with the strongest density in the A and D sites. However, the density for the drug was ill-defined (**Fig. S2a)**, not up to the level of definition expected at this resolution. We speculated that this blurring was due to imposing icosahedral averaging on a conformational ensemble. For this reason, we revisited the reconstruction process without assuming symmetry.

**Figure 3.**
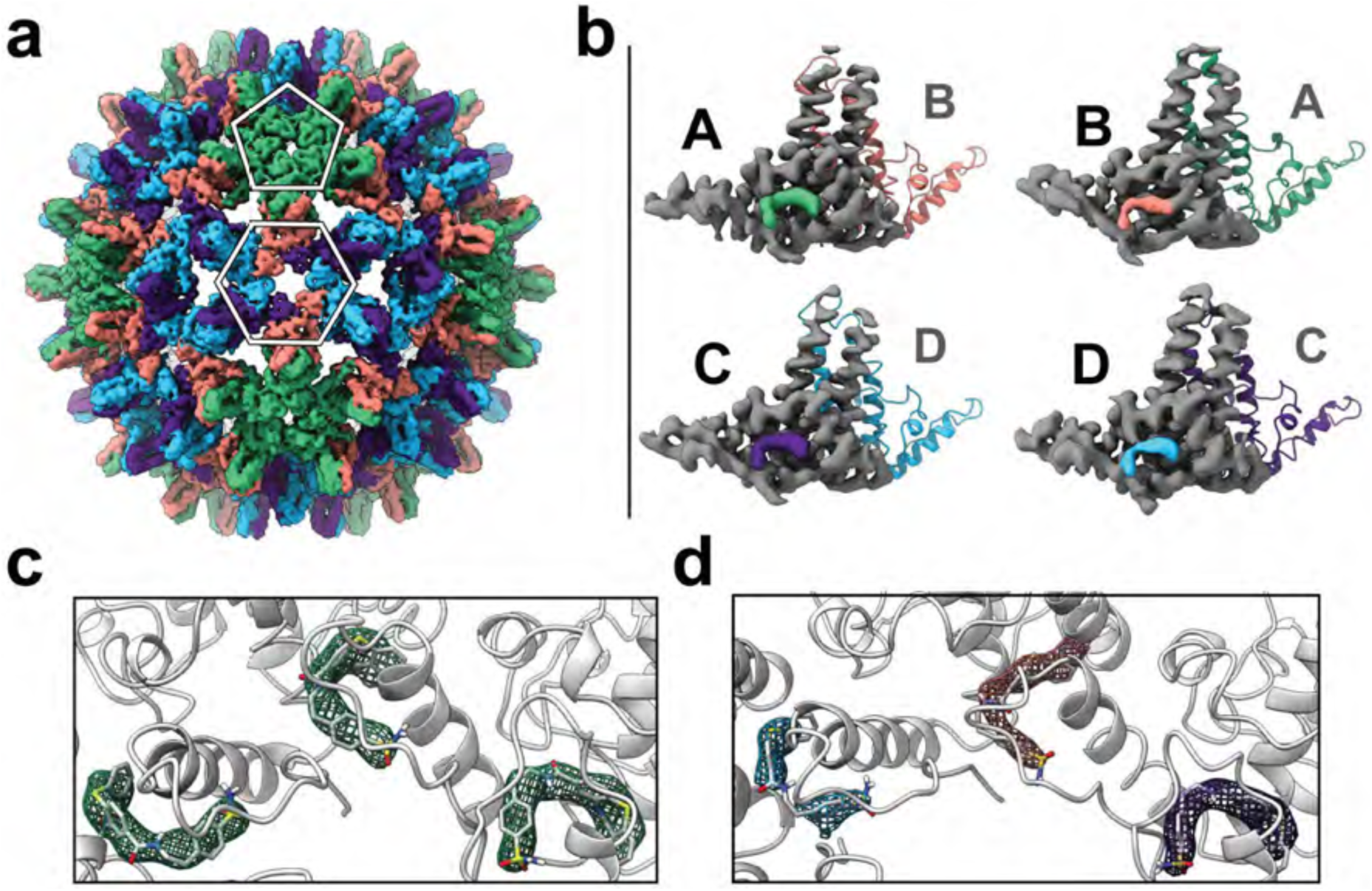
Cryo-EM reconstruction reveals density for the DBT molecule. **(a)** A T=4 HBV capsid shows the arrangement of AB dimers (green and orange, respectively) and CD dimers (purple and blue) and results four “quasi-equivalent” protein interfaces (B-C, C-D, D-B, A-A). A reconstruction of the capsid-DBT complex that employed icosahedral averaging had insufficient detail to evaluate DBT binding at every interface **(Fig. S2). (b)** Asymmetric reconstructions which focus on localized hexameric and pentameric regions of the capsid surface provide the clarity needed to identify the DBT molecule in all four sites quasi-equivalent sites (A,B,C,D) without any ambiguity. Protein density (gray) and DBT1 density (colored as in panel a) for each class of dimer is shown. **(c, d)** The DBT1 molecule binds each site in a similar pose for all sites in either the pentamer focused reconstruction **(c)** or the hexamer reconstruction **(d).** The DBT1 wraps around helix 5 of the adjacent, capping subunit with the sulfonamide group solvent-exposed towards the capsid lumen.

After performing focused asymmetric reconstruction, using either a fivefold or a quasi-sixfold as the basis structure, DBT1 structural detail was enhanced and its locations were unambiguous **(Fig. S2b)**. The CpAM density appeared as a bent plane that was consistent with the DBT1 structure **(Fig. 1a inset** and **3b,c,d).** The DBT1 density fits into the CpAM pocket formed by helix 3 (residues 26-42), helix 5 (residues 112-128), and the extended structure following helix 5 (residues 133-140). DBT1 density also wraps around helix 5 from the neighboring “capping” subunit, centered around residue 124. We find a DBT1 molecule in every quasi-equivalent protein-protein interface. This presents a very different distribution than seen with previous CpAM structures, where only B and C of the four quasi-equivalent sites were filled [30, 46., 47]. The pockets of the four quasi-equivalent sites vary substantially **(Fig. S3).** In capsids without a CpAM, the available volume at the A and D interfaces is much more restricted than in the B and C sites. However, DBT1 has the flexibility to fit into the spatially restricted sites, and consequently it has a direct effect on every dimer-dimer interaction in the capsid.

Bound CpAM affects capsid quaternary structure, typically expanding the capsid diameter by about 5%. For structural comparison, we use capsid bound to HAP-TAMRA (6BVF [46]), in part because the binding stoichiometry was verified via the TAMRA fluorophore; also the 6BVF structure was determined by cryo-EM and is not constrained by crystal packing contacts. In 6BVF, HAP-TAMRA fills the B and C sites resulting in flattening of the quasi-sixfold vertex with attendant flexing of the fivefold (where HAPs do not bind). These concerted motions make the capsid appear more faceted than a typical apo-capsid (3J2V). In comparison, the icosahedrally averaged DBT1 capsid appears to have local distortion that is intermediate between the apo-capsid and the HAP-TAMRA capsid. The presence of DBT1 at the fivefold limits the flexing of the vertex and, similarly, the loose fits of the DBT1 in the two B and two C sites do not require the quasi-sixfold to be flat. However, the fact that focused reconstruction was required to visualize details of the DBT1 structure indicates that to maintain local structure in most vertices, capsids were asymmetric and may include a subset of capsomers that were locally deformed to absorb defects over the rest of the capsid.

To investigate the difference in quasi-equivalent specificity between DBT1 and HAPs, we docked the HAP molecule NVR10-001E2, from a relatively high resolution structure [48], into the DBT1 structure (**Fig. 4**). We find there are major clashes that would not permit binding without large scale movements of the capping dimer **(Fig. 4c,e**). Residue V124 especially, is not visible in panels **c** and **e** because it is completely overlapped by the HAP. In contrast, DBT1 avoids such a clash because of its flexible chemical structure, twisting around the capping helix to avoid V124 **(Fig. 4d,f).** Other serious clashes which were visible when docking the HAP into the A or D sites include I105 and I109, which have been identified as sites of HAP-resistant mutants [48]; I105 and I109 help occlude the A and D sites from HAPs and presumably confer resistance by interfering with HAP entry to the B and C sites [49]. These clashes highlight the importance of molecular scaffold as a determinant of site specificity. The V124 clash is with the core of the HAP molecule, not with a substituent. Thus, it seems unlikely that any HAP derivative could be synthesized which binds in the same manner as DBT. Even though both molecules bind the same protein interface, the specifics of their molecular structure matter, and lead to different structural properties and assembly phenotypes.

**Figure 4.**
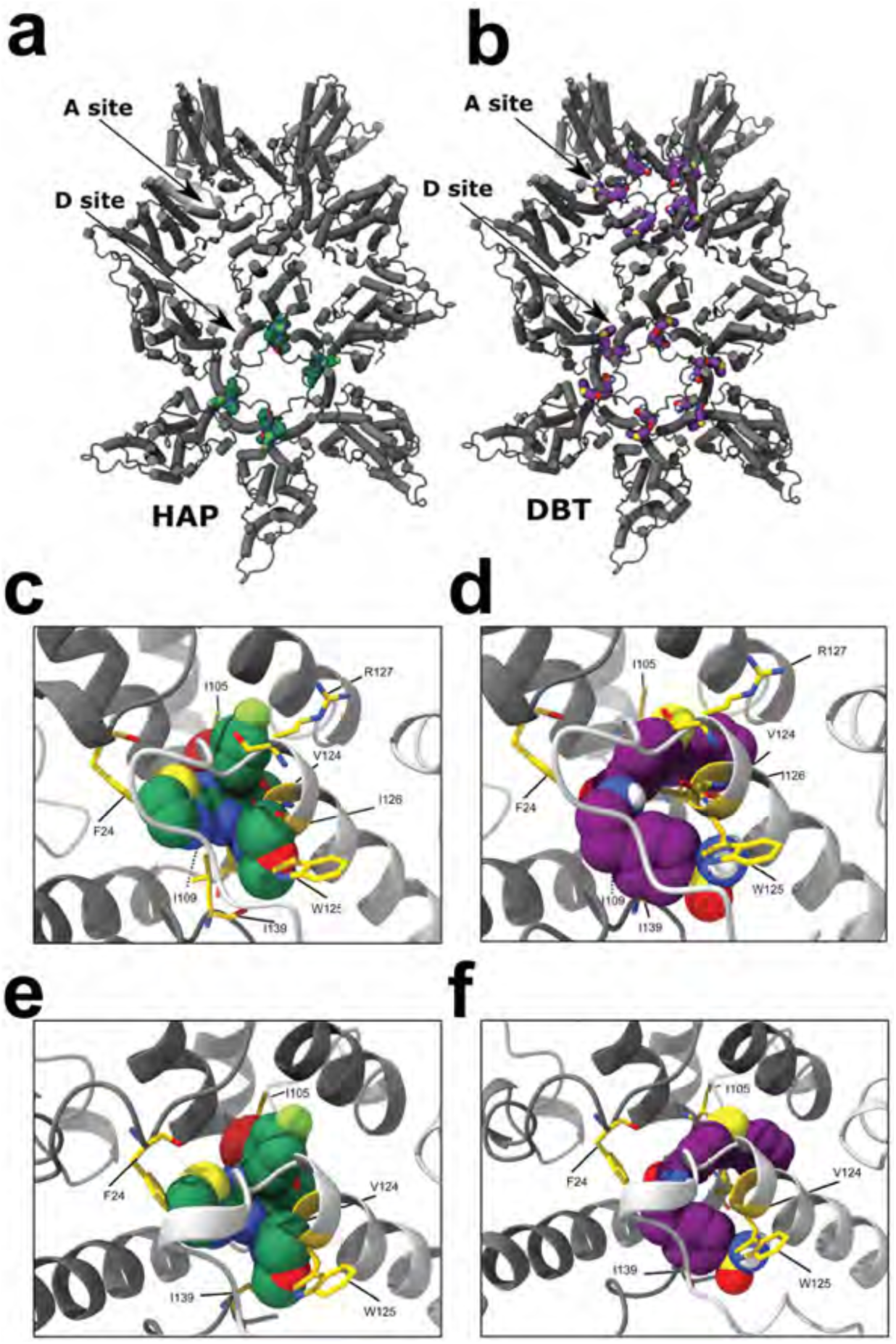
DBT maximizes occupancy of sites on the capsid surface. **(a)** Other CpAMs will only bind a subset of available interfaces. HAPs (and PPAs) bind to the B and C sites. **(b)** DBT1 also binds A and D sites. **(c)** We docked a HAP1 molecule [40] into the A site of the DBT structure and observe steric clashes. **(c: clashing residues in yellow). (d)** For comparison, DBT1 fits the site without collisions. **(e, f)** The same exercise was performed for the D site. Mutations to several of the clashing residues are known to confer resistance to HAP molecules in the B and C sites. When DBT binds these sites (d,f) there are not any clashes. The DBT molecule twists around the capping dimer, making continuous contact with V124 from the capping subunit.

To compare how multiple CpAM chemotypes bind the same pocket, in (**Fig. 5)** we summarize the binding poses of unique chemotypes of known structure. The degeneracy of the interaction between core protein and CpAMs is striking; a single CpAM will bind to multiple quasi-equivalent interfaces, and each interface will bind multiple CpAM chemotypes. With DBT1 we introduce a molecule which maximizes degeneracy by binding every quasi-equivalent site on the capsid surface. Also unique among these chemotypes, is the ability of DBT1 to engage the neighboring subunit.

**Figure 5.**
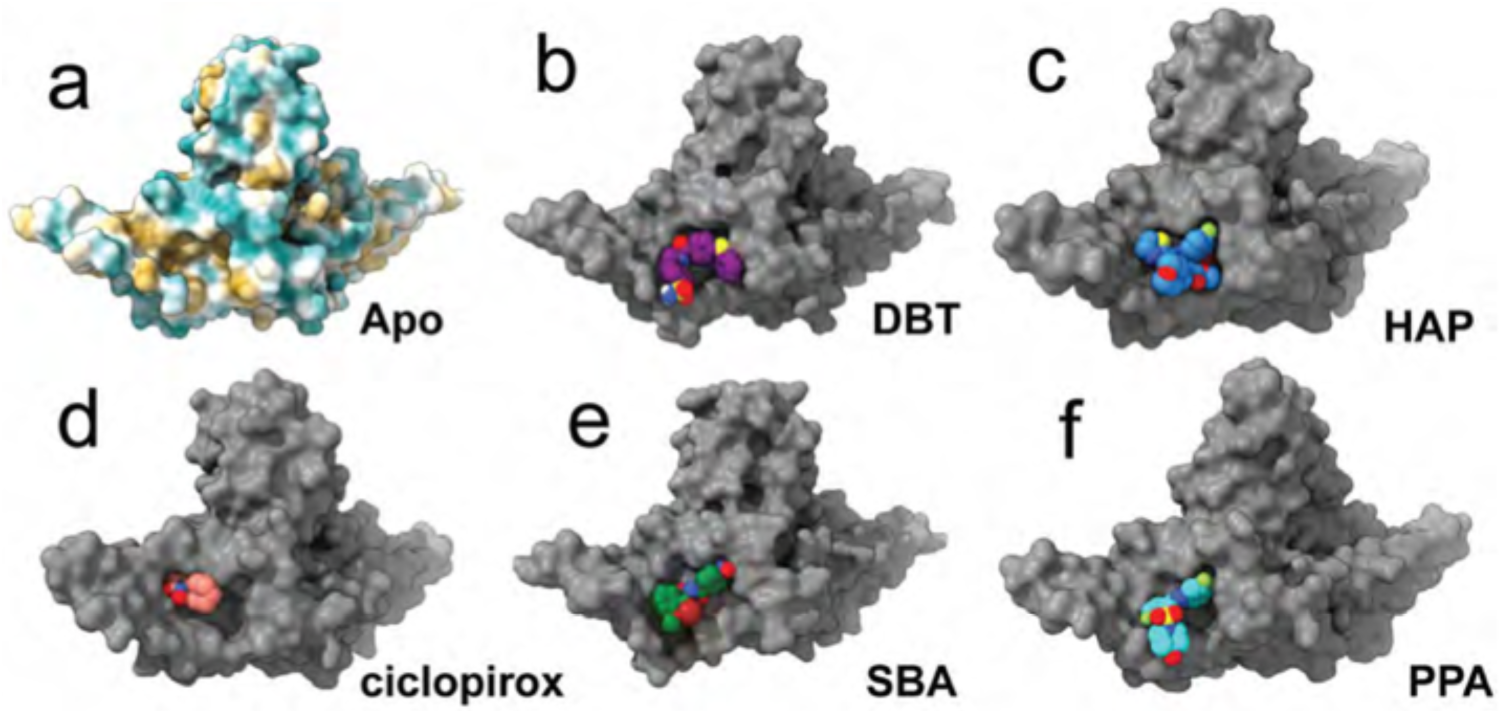
DBT1 adopts a distinct binding pose. For clarity only one half of the interface is shown, with the CpAM binding pocket visible as a cavity in the subunit surface. **(a)** The interactions at subunit interfaces are dominated by hydrophobic contacts (tan is more hydrophobic). There is a remarkable diversity of molecular scaffold for molecules which bind the inter-dimer interface. Shown are dibenzothiazepine **(DBT, b)**, heteroaryldihydropyrimidine **(HAP, c)**, ciclopirox **(d)**, sulfamoylbenzamide **(SBA, e)** and phenylpropenamide **(PPA, f)**.

## Discussion

The virus capsid as a target for therapeutics has some important advantages over other targets. Most viral structural proteins have little similarity to host proteins. Polymeric complexes, like capsids, may suppress the effect of resistant mutants, because the presence of any wildtype protein will exert a dominant sensitive effect [50]. CpAMs directed against the HBV capsid are assembly agonists. HBV is not the only icosahedral virus that has been targeted. A series of anti-picornavirus molecules stabilized the capsids to inhibit genome release [51]. A series of HIV maturation inhibitors prevent proteolysis of the Gag polyprotein presumably by stabilizing Gag-Gag interactions [52]. These capsid-directed molecules either are or were in clinical trials. All of them are capsid stabilizing. This makes sense from a physical chemical perspective, because assembly is reaction that has a steep energetically downhill [29, 53]. These molecules take advantage of mechanisms that nature has already established.

In this paper we show that DBT1 is an assembly agonist with a range of functions. DBT1 stabilizes each protein-protein interaction by approximately 1.3 kcal/mol; this effect is amplified because each dimer is tetravalent, and the resulting capsid has 240 protein contacts. Because stronger association energy stabilizes intermediates, this thermodynamic effect contributes to faster assembly kinetics [54-58]. At a structural level, DBT1 has a unique pose in the CpAM pocket, requiring less volume and interacting more strongly with helix 5 of the capping subunit. DBT1 also fits into all four quasi-equivalent sites of a T=4 particle, a promiscuity never previously seen. Depending on conditions, DBT1 can lead to non-capsid polymers indicating a competition between normative and aberrant assembly paths [59]. Accelerated assembly and mis-assembly are thus a first basis of antiviral activity. The ability of CpAMs like DBT1 to promote assembly in the absence of the pgRNA-polymerase complex indicates nucleation as a second basis for accelerated kinetics and antiviral activity [36]. Paradoxically, we also show that DBT1 can destabilize capsids leading to their dissociation, an effect that may be of particular importance for antiviral effect on metastable, mature, rcDNA-filled capsids [44, 45].

We have summarized Cp and CpAM oligomerization as equilibrium reactions and a free energy diagram **(Fig. 6)**. Starting with free subunit in the absence of any CpAM, assembly will produce normal icosahedral capsids **(1a).** This process is thought to be initiated by Cp adopting an assembly-active state and forming a nucleating complex, which may be a trimer of dimers [60-63]. Because each incoming subunit during assembly makes progressively more contacts, the assembly energy surface has a progressively steeper down-hill slope [64]. When a CpAM is added to pre-formed capsids, it will bind the capsid **(1b)**. By thermodynamic linkage [65], this necessarily lowers capsid energy. Because CpAM binding modifies the inter-subunit interaction, the bound capsid will be in a “strained” state. In free energy terms, the capsid takes on a global penalty, a positive free energy (ΔG_strain_), which is paid for in part by CpAMs having a high affinity for the local subunit interface. If the strain penalty is sufficiently large, the capsid may be able to reach a lower free energy through rupture **(1c)**. Though the rupture event breaks some local contacts, it relieves the global strain penalty. Formally, the free energy of a strained capsid is the sum of its local contact terms, minus a global strain penalty:

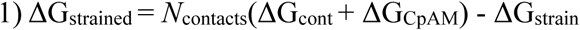

**Figure 6.**
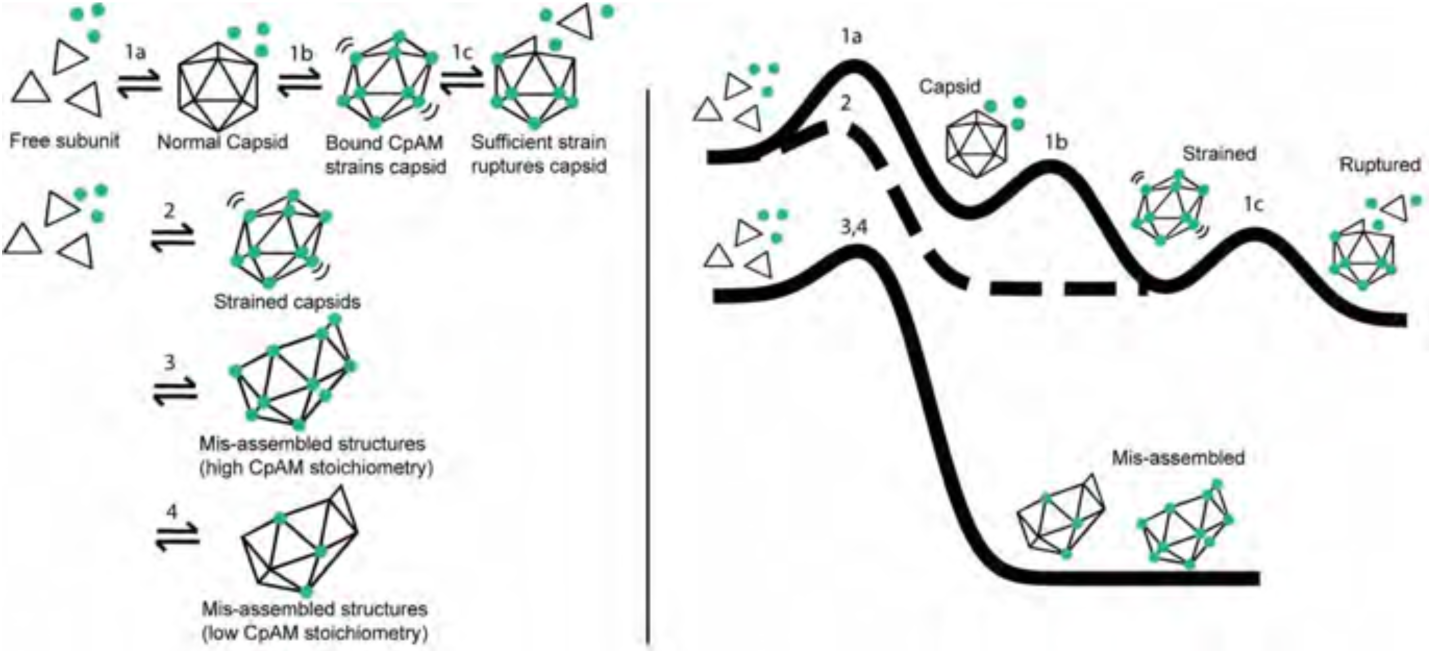
Free energy schematic for CpAM mechanisms of action. On the left side we describe a series of reactions, on the right side we describe free energy diagrams for these reactions**. (Reaction 1)** In **1a** free subunit assembles into a normal capsid. In **1b** CpAM binds to the pre-assembled capsid. We know this is a lower free energy state, because the CpAM binds. We assume all bound capsids are globally strained, because CpAMs bind to subunit interfaces, and we observed PPAs, which don’t cause aberrant assembly, cause capsids to expand by 5%. In **1c**, if the strain is sufficient the capsid ruptures releasing free subunit. **(Reaction 2)** Capsid and CpAM co-assemble to form capsids. These capsids assemble into icosahedral, or nearly icosahedral particles, but because they have bound CpAM they are strained in that they are conformationally and energetically distinct from the apo capsid**. (Reaction 3)** Co-assembly of free subunit and CpAM lead to aberrant mis-assembled products. We depict this reaction with saturating CpAM, with one drug at every subunit interface. For this reaction, a CpAM at every subunit interface is required for mis-assembly (as seen with BAY 41-4109 [43]). **(Reaction 4)** Assembly produces similar morphology to reaction 3, but at sub-stoichiometric CpAM. This implies kinetic regulation, perhaps at the level of nucleation, leading to aberrant assembly. Free subunit from ruptured capsid (product of reaction **1c**), may enter reactions **3** or **4** to form aberrant structures.

Where the subunits in a capsid are connected by *N*_contacts_, each with and association energy of ΔG_cont_; the bound CpAM contributes ΔG_CpAM_ to this association energy. For Cp149 and DBT1, ΔG_cont_ is −3.1 Kcal/mol/cont and ΔG_CpAM_ is −1.3 kcal/mol/cont (**Fig. 1b inset**). The free energy of a ruptured capsid has fewer local contacts, but less global strain penalty:

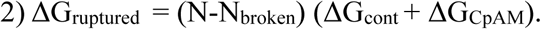

While ruptured capsids can reanneal, if ΔG_ruptured_ < ΔG_strained_ they will preferentially remain in a ruptured state and possibly release free subunits. The extent of global strain which can be tolerated will be dependent on the affinity of the CpAM for the local interfaces, the energy increment to the interface from the CpAM (ΔG_CpAM_) and ΔG_cont_. For HAP-TAMRA we were able to directly observe the strained state at equilibrium [66]. For the more potent HAP12, we often see the larger misassembled products (**Fig 2a,f**) that are the ultimate products of freed subunits. DBT1 was the only CpAM we tested where we could observe released free subunit (**Fig. 2a,b**). In comparison, the SBA that we tested showed no evidence of capsid rupture, indicating that some CpAMs induce a modest strain penalty (ΔG_ruptured_ >> ΔG_strained_).

Co-assembly of capsids from subunits with CpAM can lead to approximately normal and clearly abnormal assembly products, depending on the CpAM and reaction conditions (**Fig. 6, Reactions 2-4**). Assembly with an SBA or PPA “always” leads to normal capsids (**Fig 6. Reaction 2**); however, a PPA-capsid crystal structure shows that the CpAM results in a capsid that has a radius that is 5% larger [47]. Assembly with sub-stoichiometric concentrations of HAP (e.g. BAY41-4109) also lead to morphologically normal capsids [43, 67]. Conversely, assembly with super-stoichiometric concentrations of HAP leads to aberrant assembly (**Fig 6. Reaction 3**). Apparently, the HAPs tested here must be present at almost every possible site to sustain the aberrant morphology. While co-assembly with DBT1 at sub-stoichiometric concentrations preferentially forms morphologically normal capsids, it will also produce some aberrant structures (**Fig 2c**). This suggests that a CpAM can induce aberrant geometry at an early point in assembly, possibly nucleation, and that defect will persist (**Fig 6. Reaction 4**). Of note, subunits released for ruptured capsids (**Reaction 1c**) may feed into co-assembly reactions, notably reactions **3** and **4**, consistent with the appearance of aberrant capsids after incubation of normal capsids with HAP12. Because intact capsids, even when strained, will have to break many intersubunit contacts to rupture, there is likely to be a high energy barrier, resulting in metastable particles. On the other hand, aberrant particles always have exposed ends, allowing ready release and re-binding of subunits; thus, the presence of aberrant particles in the presence of CpAM indicates that these are a lowest energy state (**Fig. 6**).

In a biological environment, where mature capsids are at vanishingly low concentration and are metastable due to the encapsidation of a dsDNA coiled spring [44, 45], the addition of a drug that favors aberrant geometry may be enough to cause capsid rupture to disrupt the normal path of infection. (We note that a molecule that strongly stabilizes capsid may also disrupt normal genome release, as with the picornavirus-stabilizing compounds [51]) While energy diagrams are instructive, they fail to show the full complexity of CpAM behavior, even when the complexity has been reduced by using purified protein. For example, the effect of a CpAM on association energy is not fully predictive of a reduction in secreted virus from virus-expressing cells [40]. Nonetheless, association energy does explain how different CpAMs can overcome CpAM-resistant mutations [49]. Conversely, the effects of nucleic acid on assembly are unexplored. The advantage of an energy diagram for assembly suggests testable mechanisms for the effects of a CpAM on a capsid [46]. But energy diagrams are incomplete. CpAM activities may also include interfering with the ability of the capsid to support reverse transcription [20] and affecting the hypothesized roles of Cp on RNA transport [16]. For all of these reasons, we argue that categorizing CpAMs as Type 1 (aberrant assembly) or Type 2 (normal capsid morphology) are over-simplified and direct attention away from determining what mechanisms are biologically critical.

We show that DBT1 has distinct behavior in capsid assembly and destabilization relative to other CpAMs. The experiments presented here demonstrate that DBT1 paradoxically induces assembly and mis-assembly while simultaneously globally destabilizing capsid structure. There can also be significant variations within a class, depending on specific substituents. We expect that development of the most effective CpAM variants will favor molecules which make core protein dysfunctional during *multiple* processes of the viral lifecycle – capsid assembly, packaging pgRNA, mediating reverse transcription, delivering DNA to the nucleus, and maintaining the critical pool of cccDNA which sustains chronic infection – towards the ultimate goal of achieving a functional cure for patients living with chronic HBV.

## Materials and Methods

### Preparation of material

The Hepatitis B subtype *adyw* core protein assembly domain was expressed in *E. coli* using two constructs: Cp149 which is a C-terminal truncation of the core protein, and is sufficient to assemble capsids *in vitro*, and Cp150 which is a further modification of Cp149 with three native cysteines mutated to alanine and an additional C-terminal cysteine appended to the C-terminus. Both constructs were purified by ammonium sulfate precipitation followed by size exclusion chromatography, as previously described [68]. To prepare purified empty capsids, core protein homodimers were induced to assemble using 300mM NaCl, 20 mM Tris, pH=7.5, overnight. The assembled capsids are subsequently purified from un-assembled dimer subunits using a Superose 6 10/300 GL size exclusion column (GE Healthcare). The small molecule allosteric modulators were purchased (Chem-Div, San Diego).

### Quantification of assembly reactions

For the assembly reactions involving Cp149 and DBT1, fresh dimer was added at a 1:1 ratio by volume to a premixed reactant mixture of NaCl, buffer, and DBT1, so that the post-mixing concentrations were 5µM dimer, and each DBT1 concentration specified. Assembly kinetics were monitored by 90 ° light scattering using a Photon Technology fluorometer with excitation and emission monochromators at 320 nm. The mass fractions of assembled and un-assembled subunit were resolved by size exclusion chromatography using a Superose 6 column and quantified on the basis of absorbance at 280 nm.

### Detection of CpAM-induced capsid disruption

Serial dilution of DBT-1 with DMSO was performed to produce drug stocks at 0, 125 µM, 250 µM, 500 µM, 1 mM, and 2 mM concentration. 2 μL of each stock was added to 98 μL each of 150 mM, 300 mM, 600 mM, and 1,200 mM NaCl in 20 mM Tris (pH=7.5). These were then mixed at a 1:1 ratio with 10 µM Cp149 purified capsids (20 mM Tris, pH=7.5), for a final concentration of 5µM Cp149 dimer, in capsid form (in 1% DMSO). Samples were given 12 hours to incubate, before analysis. Following incubation, samples were loaded in a 1% native agarose gel in TBE running buffer for electrophoresis at 50 volts for 4 hours, then blotted for 8 hours on a polyvinylidene membrane (Immoblion-P, Millipore). The membrane was incubated with Rabbit anti-Cp primary antibody (prepared using antigen from the Zlotnick lab), then horseradish peroxidase-labeled secondary antibody (1:5000, ThermoFisher Scientific). Luminol and hydrogen peroxide in a 1:1 ratio (Thermo Scientific) were added to the membrane for imaging. To verify the results of the gel electrophoresis for DBT1, the reactions were analyzed by size exclusion chromatography on a Superose 6 10/300 GL column, using the same buffer as used for gel electrophoresis (TBE), and at a low flow rate (0.125ml/min) for comparable running times between the electrophoresis and chromatography.

### Negative stain electron microscopy

Samples were adsorbed to the glow discharged carbon film coated 300 mesh coper grids (Electron Microscopy Sciences), washed with water, stained and air dried. Samples were stained with 2% uranyl acetate **(Fig. 1)** or 6% ammonium molybdate (6%) and trehalose(0.5%) **(Fig. 2).** Trehalose is added to minimize the collapse of particles that occurs on the grid upon dehydration. Images were collected with a JEOL TEM 1010 **(Fig. 1)** or a JEOL 1400plus microscope **(Fig. 2).**

### Cryo-electron microscopy and image processing

For single particle Cryo-EM, the Cp150 construct was used to prevent capsids from dissociating before imaging. The Cp150 capsids are prepared in the same way as Cp149 capsids, but spontaneously form an inter-dimer disulfide crosslink. Purified Cp150 capsids were then incubated with a molar excess of DBT1 overnight before concentration to 10 mg/ml using an Amicon centrifugal concentrator. Concentrated samples were applied to a glow-discharged Quantifoil holey-carbon grids (R2/2). The grids were blotted with filter paper for 4 s before automated plunging into liquid ethane using an FEI vitrobot. Imaging was carried out on a FEI Titan Krios operated at 300 kV at a nominal magnification of 22,500. Images were recorded on a Gatan K2 Summit detector operating in super-resolution mode, resulting in a pixel size of 0.65 Å with a dose of ∼33 e^-^ Å^2^. Each exposure was 8 s long and was collected as 35 individual frames. Cryo-EM classification and reconstruction were implemented by adapting standard protocols of the EMAN2, and Relion software programs [69-71]. The concept of using focused sub-particle reconstruction was inspired by previous descriptions from Ilca, Huskonien et al [72, 73].

### Model Determination and Structural analysis

The crystal structures of both apo capsid (1qgt) [74] and HAP-TAMRA bound capsid (6bvf) [66] were used as starting points for flexible model refinement using the PHENIX, COOT, and eLBOW software programs.[75-77] The model validation statistics we report were obtained from the final output of the Phenix real space refinement tool. Structural comparisons, figure creation, and molecular docking tasks were carried out in UCSF Chimera and ChimeraX [78, 79].

## Acknowledgements

This effort was funded by NIH grant R01-AI144022 to AZ. Some EM work was supported by a grant from the Indiana Clinical and Translational Sciences Institute (CTSI) to AZ. We made use of the Indiana University Electron Microscopy Center, the Purdue University Cryo-EM center, and the IU Physical Biochemistry Instrumentation facility. AZ has an interest in biotech companies developing HBV-directed antiviral agents. SD is now an employee of Door Pharmaceuticals.

## Supplementary Figures

**Figure S1.**
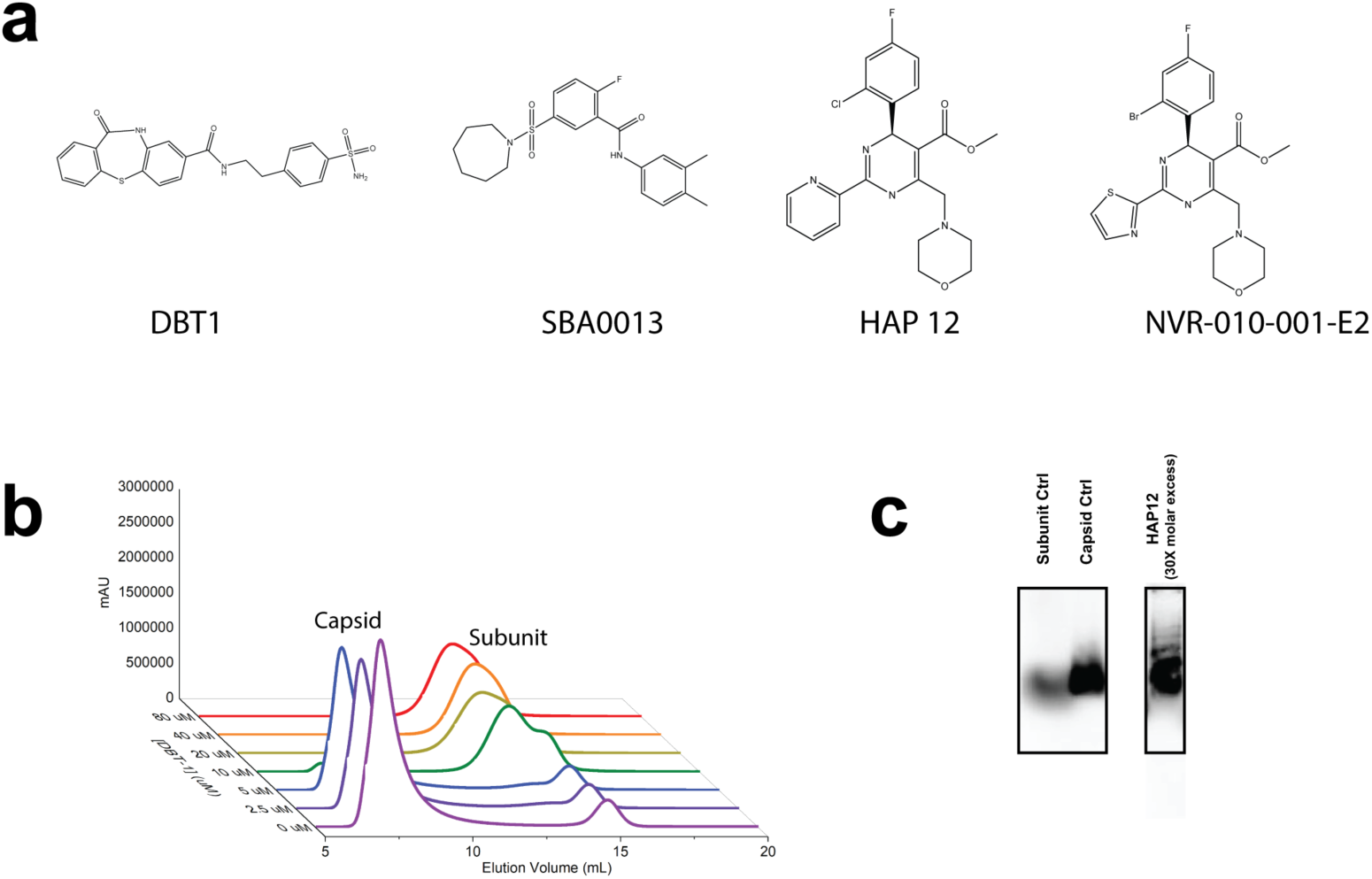
Supplemental chemical structures, size exclusion chromatography, and agarose gel. **(a**) Each of the chemical structures used experimentally (DBT1, SBA0013, and HAP12) along with the HAP molecule NVR-010-001-E2, which derives from the high-resolution crystal structure 5E0I, and was used for the molecular docking in (**Fig. 4).** Panel **(b)** uses SEC to recapitulate our observations from the agarose gel electrophoresis in **(Fig. 2a,b)**, that DBT1 causes capsids to transition to free subunit. Using shorter running times for the SEC did not detect an increasing dimer pool, consistent with the EM results in **(Fig. 2d).** For us this makes a compelling kinetic argument, that the broken capsid state only exists transiently when treated with DBT1. To further emphasize the disruptive effects of HAP12 on pre-formed capsids, we include another agarose gel blot **(c)**. Here the agarose gel percentage was higher (1.2%) than in **Fig. 2a** (1%), and the distribution of larger mis-assembled products becomes visible. However, the resolution between subunit and capsid become poor, so we opted for the lower agarose condition (1%) for subsequent experiments.

**Figure S2.**
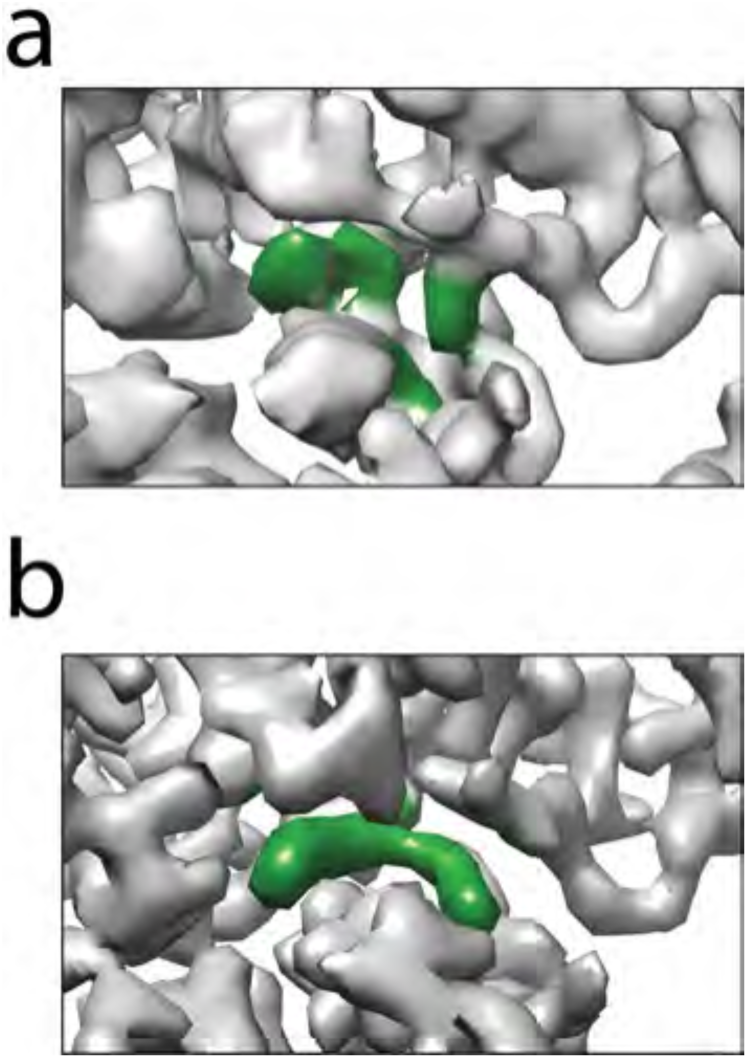
Focused reconstruction reveals detail unseen when imposing icosahedral symmetry. Colored in green is the density in the CpAM pocket. Our initial reconstruction, imposing icosahedral symmetry, had weak and incoherent density for the DBT1 molecule **(a)**. An alternative reconstruction approach, focused asymmetric reconstruction **(b)**, does not assume perfect capsid symmetry, and dramatically improves the assignment of DBT1 drug density. It is important to note here that both reconstructions depicted here are binned by 2X and have not been processed to the maximum extent supported in Relion (no unbinned processing, no CTF refinement, no Bayesian polishing). Once we saw the dramatic improvement visible here, we abandoned the icosahedral refinement. Thus, it is possible that further icosahedral processing would indeed resolve drug density, but we expect the detail always be worse than what could be achieved with a focused subparticle treatment. This is due to structural variability, both inherent to the capsids and variability imposed by the DBT1-induced strain.

**Figure S3.**
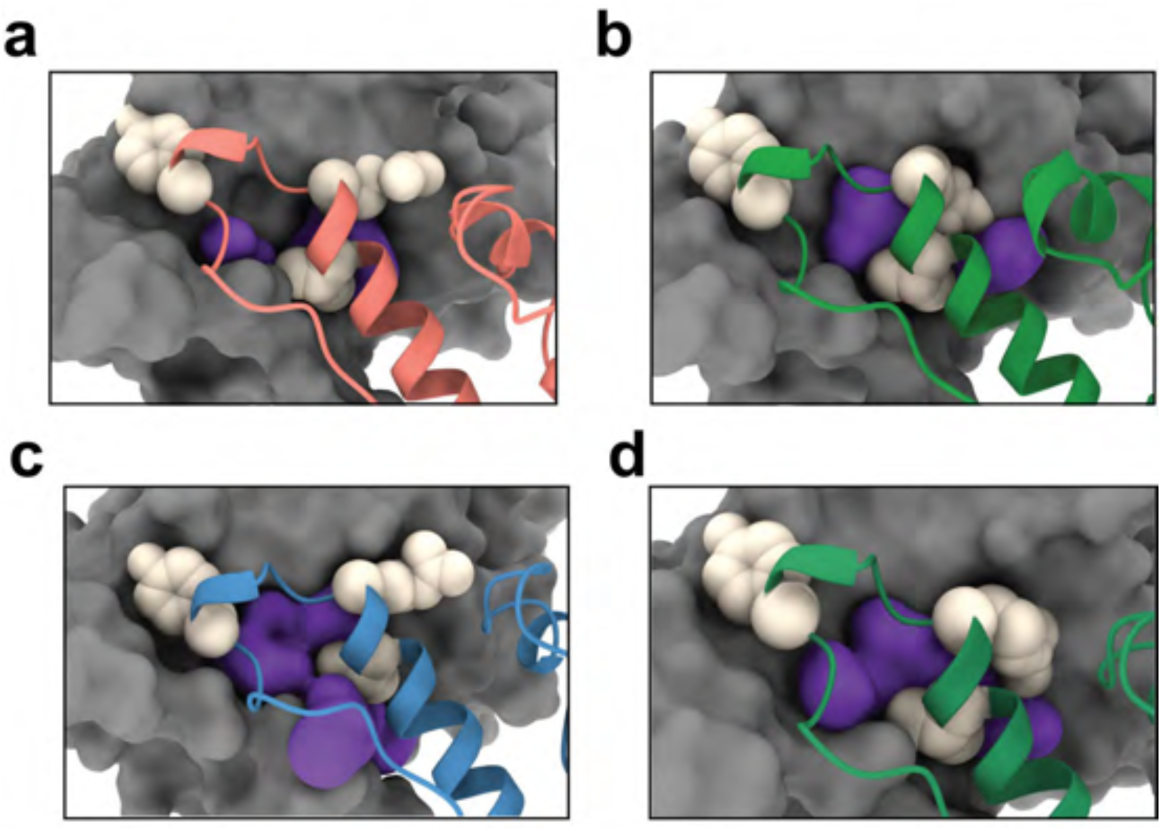
Quasi-equivalence and interface volume. A view of each of the four quasi-equivalent interfaces on the T=4 apo cryo-EM structure (3j2v). Void space is rendered as a purple surface, with important interface residues shown in white (Y132, R127, V124). Despite being composed of identical protein surfaces, each interface is slightly different. This demonstrates the principle of quasi-equivalence and explains how CpAM molecules might bind each interface in a different manner. Interface A **(a)** has basically zero available space in the apo structure. For DBT to bind this site necessitates allosteric changes in quaternary structure (See **Fig. S4).** Site D **(b)** is also quite restrictive, and neither of these sites have been observed to bind HAPs. In comparison, sites B **(c)** and C **(d)** have considerably more accessible volume in the apo capsid, which offers one explanation for how HAPs can bind to these sites.

**Figure S4.**
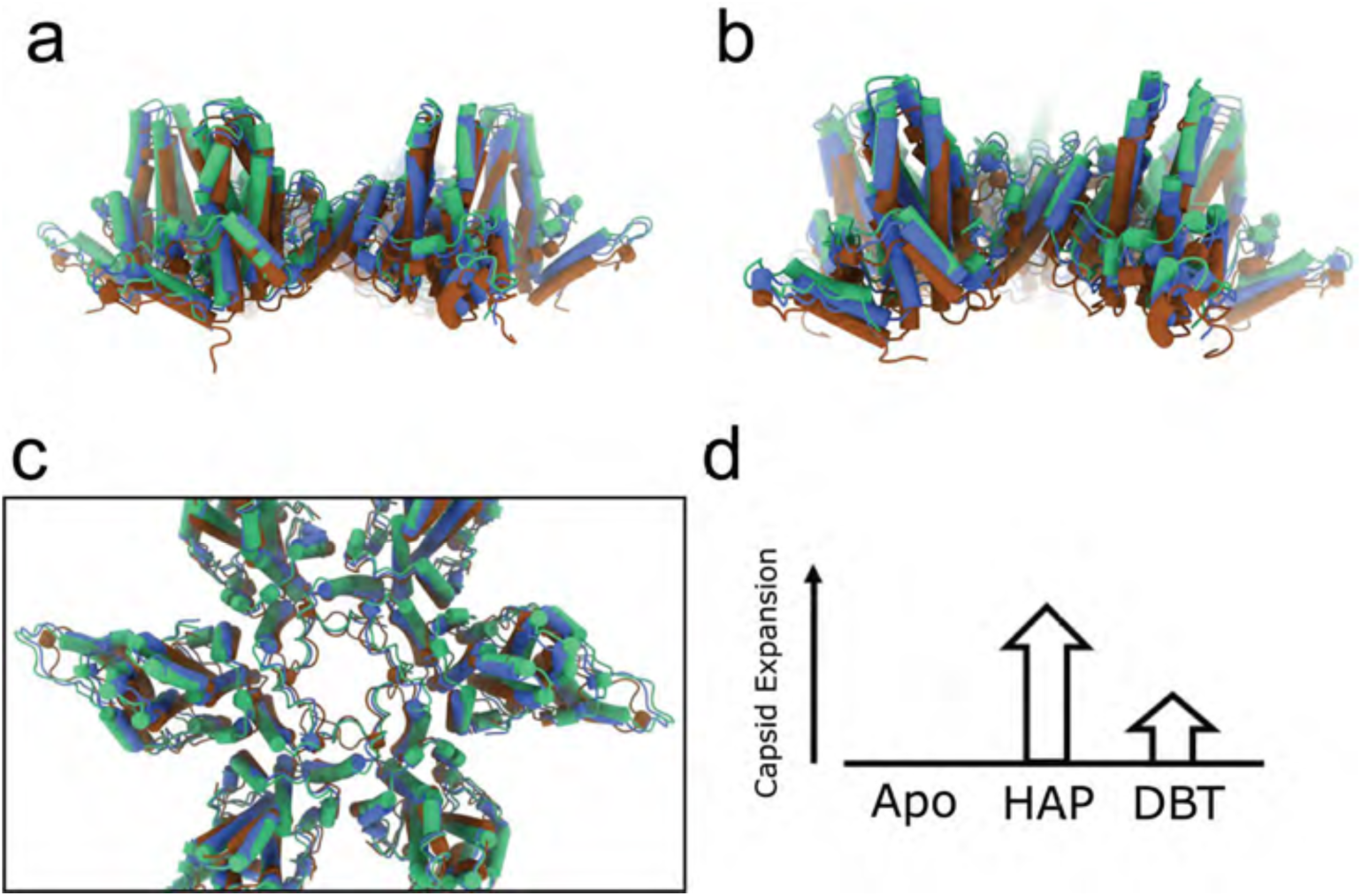
Changes in quaternary structure upon DBT binding. The DBT model (blue) is compared to two references: apo capsid (3J2V, brown) and HAP bound (6BVf, green). Side views of the hexamer **(a)** and pentamer **(b)** demonstrate the variation in capsid diameter between the three structures. Similarly, conclusions are drawn from a top view of the hexamer **(c)**: the structures are similar but contain differences in quaternary structure. Because of the oligomeric nature of the capsid surface, these differences become amplified the further away one compares from the point of alignment. Here the point of alignment is the center of the capsid, thereby equalizing structural differences across the capsid. A schematic of the change in capsid expansion is shown in **(d)**, where the DBT causes a return towards the apo form relative to the HAP bound capsid, despite containing more drug molecules.

**Figure S5.**
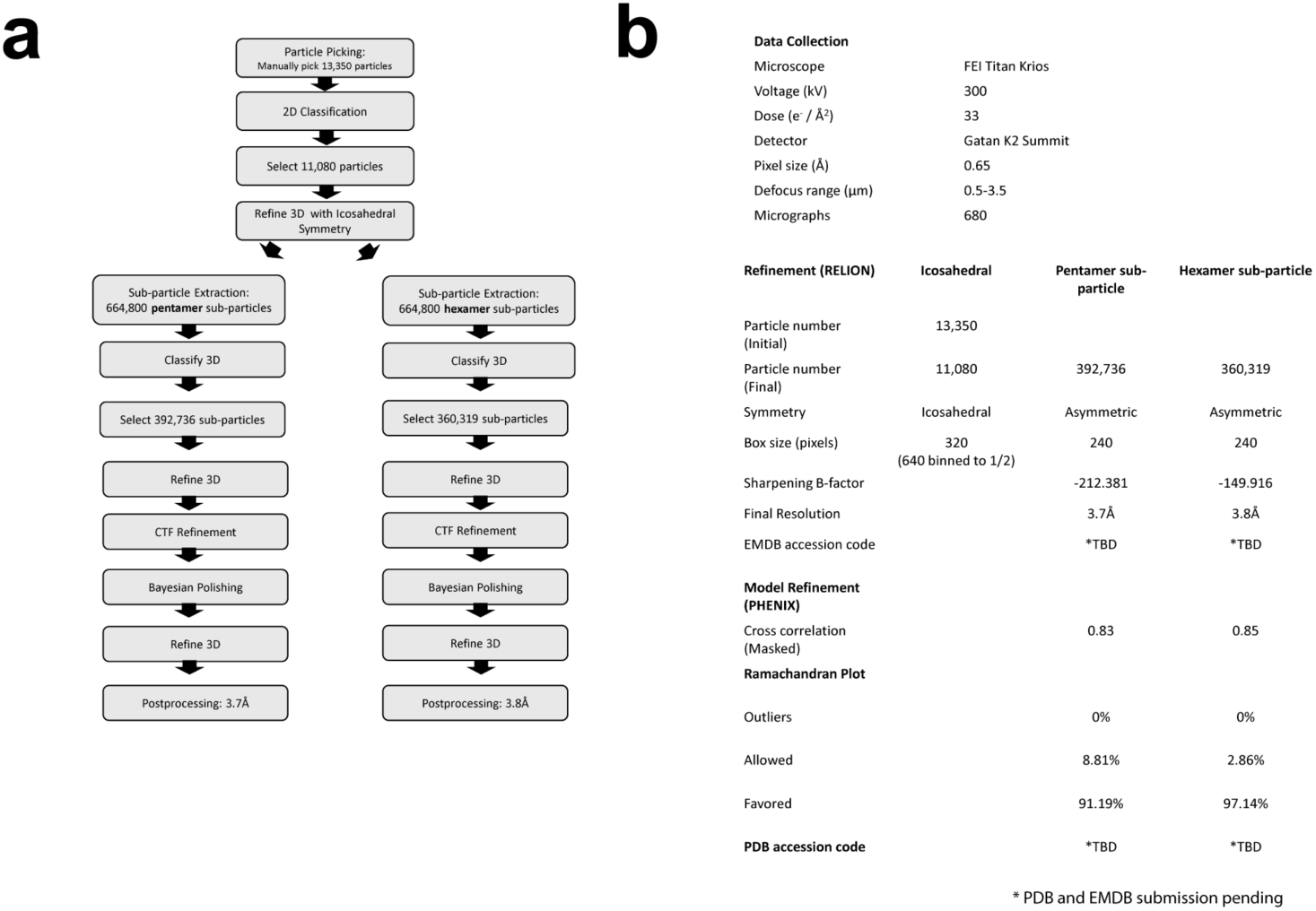
Cryo-EM reconstruction details. A flowchart **(a)** outlines the reconstruction process we used for the cryo-EM image reconstruction. All major steps were performed using Relion and EMAN2 [69-71]. Statistics which describe the microscopy, data processing, and model fitting are presented in table **(b).**

## Bibliography

1. Polaris Observatory, C., Global prevalence, treatment, and prevention of hepatitis B virus infection in 2016: a modelling study. Lancet Gastroenterol Hepatol, 2018. 3(6): p. 383–403.

2. Graber-Stiehl, I., The silent epidemic killing more people than HIV, malaria or TB. Nature, 2018. 564: p. 24–26.

3. Liang, T.J., Hepatitis B: the virus and disease. Hepatology (Baltimore, Md.), 2009. 49(5 Suppl): p. S13–S21.

4. Ni, Y.H., et al., Two decades of universal hepatitis B vaccination in taiwan: impact and implication for future strategies. Gastroenterology, 2007. 132(4): p. 1287–93.

5. Grossi, G., et al., Hepatitis B virus long-term impact of antiviral therapy nucleot(s)ide analogues (NUCs). Liver Int, 2017. 37 Suppl 1: p. 45–51.

6. Alter, H., et al., A research agenda for curing chronic hepatitis B virus infection. Hepatology, 2018. 67(3): p. 1127–1131.

7. Yan, H., et al., Sodium taurocholate cotransporting polypeptide is a functional receptor for human hepatitis B and D virus. eLife, 2012. 1: p. e00049.

8. Kann, M., A. Schmitz, and B. Rabe, Intracellular transport of hepatitis B virus. World J Gastroenterol, 2007. 13(1): p. 39–47.

9. Mantyla, E., et al., Viral highway to nucleus exposed by image correlation analyses. Sci Rep, 2018. 8(1): p. 1152.

10. Rabe, B., et al., Nuclear import of hepatitis B virus capsids and release of the viral genome. PNAS, 2003. 100(17): p. 9849–9854.

11. Guo, J.T. and H. Guo, Metabolism and function of hepatitis B virus cccDNA: Implications for the development of cccDNA-targeting antiviral therapeutics. Antiviral Res, 2015. 122: p. 91–100.

12. Schreiner, S. and M. Nassal, A Role for the Host DNA Damage Response in Hepatitis B Virus cccDNA Formation-and Beyond? Viruses, 2017. 9(5).

13. Bock, C.T., et al., Structural organization of the hepatitis B virus minichromosome. Journal of molecular biology, 2001. 307(1): p. 183–96.

14. Belloni, L., et al., Nuclear HBx binds the HBV minichromosome and modifies the epigenetic regulation of cccDNA function. Proceedings of the National Academy of Sciences of the United States of America, 2009. 106(47): p. 19975–9.

15. Abraham, T.M., et al., Characterization of the contribution of spliced RNAs of hepatitis B virus to DNA synthesis in transfected cultures of Huh7 and HepG2 cells. Virology, 2008. 379(1): p. 30–7.

16. Li, H.C., et al., Nuclear export and import of human hepatitis B virus capsid protein and particles. PLoS pathogens, 2010. 6(10): p. e1001162.

17. Ning, X., et al., Secretion of genome-free hepatitis B virus--single strand blocking model for virion morphogenesis of para-retrovirus. PLoS pathogens, 2011. 7(9): p. e1002255.

18. Hu, J. and K. Liu, Complete and Incomplete Hepatitis B Virus Particles: Formation, Function, and Application. Viruses, 2017. 9(3).

19. Nassal, M., Hepatitis B viruses: reverse transcription a different way. Virus Res, 2008. 134(1-2): p. 235–49.

20. Tan, Z., et al., The interface between hepatitis B virus capsid proteins affects self-assembly, pregenomic RNA packaging, and reverse transcription. J Virol, 2015. 89(6): p. 3275–84.

21. Venkatakrishnan, B. and A. Zlotnick, The Structural Biology of Hepatitis B Virus: Form and Function. Annual Reviews of Virology, 2016. 3: p. 429–451.

22. Liu, K., et al., Multiple roles of core protein linker in hepatitis B virus replication. PLoS Pathog, 2018. 14(5): p. e1007085.

23. Watts, N.R., et al., The morphogenic linker peptide of HBV capsid protein forms a mobile array on the interior surface. The EMBO journal, 2002. 21(5): p. 876–84.

24. Ceres, P. and A. Zlotnick, Weak protein-protein interactions are sufficient to drive assembly of hepatitis B virus capsids. Biochemistry, 2002. 41(39): p. 11525–31.

25. Harms, Z.D., et al., Monitoring Assembly of Virus Capsids with Nanofluidic Devices. ACS Nano, 2015. 9(9): p. 9087–96.

26. Lutomski, C.A., et al., Multiple Pathways in Capsid Assembly. J Am Chem Soc, 2018. 140(17): p. 5784–5790.

27. Lutomski, C.A., et al., Hepatitis B Virus Capsid Completion Occurs through Error Correction. J Am Chem Soc, 2017. 139: p. 16932–16938.

28. Asor, R., et al., Assembly Reactions of Hepatitis B Capsid Protein into Capsid Nanoparticles Follow a Narrow Path through a Complex Reaction Landscape. ACS Nano, 2019. 13(7): p. 7610–7626.

29. Katen, S.P. and A. Zlotnick, Thermodynamics of Virus Capsid Assembly. Methods in Enz., 2009. 455: p. 395–417.

30. Venkatakrishnan, B., et al., Hepatitis B Virus Capsids Have Diverse Structural Responses to Small-Molecule Ligands Bound to the Heteroaryldihydropyrimidine Pocket. J Virol, 2016. 90(8): p. 3994–4004.

31. Venkatakrishnan, B., et al., Structure and dynamics of adeno-associated virus serotype 1 VP1-unique N-terminal domain and its role in capsid trafficking. J Virol, 2013. 87(9): p. 4974–84.

32. Feld, J.J., et al., The phenylpropenamide derivative AT-130 blocks HBV replication at the level of viral RNA packaging. Antiviral Res, 2007. 76(2): p. 168–77.

33. King, R.W., et al., Inhibition of human hepatitis B virus replication by AT-61, a phenylpropenamide derivative, alone and in combination with (-)beta-L-2’,3’-dideoxy-3’-thiacytidine. Antimicrob Agents Chemother, 1998. 42(12): p. 3179–86.

34. Deres, K., et al., Inhibition of hepatitis B virus replication by drug-induced depletion of nucleocapsids. Science, 2003. 299(5608): p. 893–6.

35. Stoltefuss, J., et al., New dihydropyrimidine derivatives and their corresponding mesomers useful as antiviral agents. 1999, Bayer Aktiengesellschaft, Germany: WO. p. 52 pp.

36. Katen, S.P., et al., Trapping of Hepatitis B Virus capsid assembly intermediates by phenylpropenamide assembly accelerators. ACS Chem Biol, 2010. 5: p. 1125–36.

37. Stray, S.J., et al., A heteroaryldihydropyrimidine activates and can misdirect hepatitis B virus capsid assembly. Proc Natl Acad Sci U S A, 2005. 102(23): p. 8138–43.

38. Campagna, M.R., et al., Sulfamoylbenzamide derivatives inhibit the assembly of hepatitis B virus nucleocapsids. Journal of virology, 2013. 87(12): p. 6931–42.

39. Kang, J.-A., et al., Ciclopirox inhibits Hepatitis B Virus secretion by blocking capsid assembly. Nature communications, 2019. 10(1): p. 2184.

40. Bourne, C., et al., Small-Molecule Effectors of Hepatitis B Virus Capsid Assembly Give Insight into Virus Life Cycle. J Virol, 2008. 82: p. 10262–10270.

41. Turner, W.W., et al., HEPATITIS B CORE PROTEIN ALLOSTERIC MODULATORS, USPTO, Editor. 2019, IURTC: USA. p. 219.

42. Bartenschlager, R. and H. Schaller, Hepadnaviral assembly is initiated by polymerase binding to the encapsidation signal in the viral RNA genome. EMBO J., 1992. 11: p. 3413–3420.

43. Stray, S.J. and A. Zlotnick, BAY 41-4109 has multiple effects on Hepatitis B virus capsid assembly. J Mol Recognit, 2006. 19: p. 542–548.

44. Dhason, M.S., et al., Differential assembly of Hepatitis B Virus core protein on single- and double-stranded nucleic acid suggest the dsDNA-filled core is spring-loaded. Virology, 2012. 430(1): p. 20–9.

45. Cui, X., et al., Maturation-associated destabilization of hepatitis B virus nucleocapsid. J Virol, 2013. 87(21): p. 11494–503.

46. Schlicksup, C.J., et al., Hepatitis B virus core protein allosteric modulators can distort and disrupt intact capsids. Elife, 2018. 7: p. pii: e31473.

47. Katen, S.P., et al., Assembly-directed antivirals differentially bind quasiequivalent pockets to modify hepatitis B virus capsid tertiary and quaternary structure. Structure, 2013. 21(8): p. 1406–16.

48. Klumpp, K., et al., High-resolution crystal structure of a hepatitis B virus replication inhibitor bound to the viral core protein. Proc Natl Acad Sci U S A, 2015. 112: p. 15196–201.

49. Ruan, L., J.A. Hadden, and A. Zlotnick, Assembly Properties of Hepatitis B Virus Core Protein Mutants Correlate with Their Resistance to Assembly-Directed Antivirals. Journal of Virology, 2018. 92(20).

50. Tanner, E.J., et al., Dominant drug targets suppress the emergence of antiviral resistance. Elife, 2014. 3.

51. Tsang, S.K., et al., Stabilization of poliovirus by capsid-binding antiviral drugs is due to entropic effects. J Mol Biol, 2000. 296(2): p. 335–40.

52. Wang, M., et al., Quenching protein dynamics interferes with HIV capsid maturation. Nat Commun, 2017. 8(1): p. 1779.

53. Perlmutter, J.D. and M.F. Hagan, Mechanisms of virus assembly. Annu Rev Phys Chem, 2015. 66: p. 217–39.

54. Hagan, M.F., O.M. Elrad, and R.L. Jack, Mechanisms of kinetic trapping in self-assembly and phase transformation. J Chem Phys, 2011. 135(10): p. 104115.

55. Michaels, T.C.T., et al., Kinetic constraints on self-assembly into closed supramolecular structures. Sci Rep, 2017. 7(1): p. 12295.

56. Endres, D. and A. Zlotnick, Model-based Analysis of Assembly Kinetics for Virus Capsids or Other Spherical Polymers. Biophys. J., 2002. 83: p. 1217–1230.

57. Porterfield, J.Z. and A. Zlotnick, An Overview of Capsid Assembly Kinetics, in Emerging Topics in Physical Virology, P.G. Stockley and R. Twarock, Editors. 2010, Imperial College Press: London.

58. Selzer, L., S.P. Katen, and A. Zlotnick, The hepatitis B virus core protein intradimer interface modulates capsid assembly and stability. Biochemistry, 2014. 53(34): p. 5496–504.

59. Kondylis, P., et al., Competition between Normative and Drug-Induced Virus Self-Assembly Observed with Single-Particle Methods. J Am Chem Soc, 2018.

60. Zhao, Z., et al., Structural Differences between the Woodchuck Hepatitis Virus Core Protein in the Dimer and Capsid States Are Consistent with Entropic and Conformational Regulation of Assembly. Journal of Virology, 2019. 93(14): p. pii: e00141-19.

61. Zlotnick, A., et al., A theoretical model successfully identifies features of hepatitis B virus capsid assembly. Biochemistry, 1999. 38(44): p. 14644–14652.

62. Packianathan, C., et al., Conformational changes in the Hepatitis B virus core protein are consistent with a role for allostery in virus assembly. J Virol, 2010. 84: p. 1607–15.

63. Zandi, R., et al., Classical nucleation theory of virus capsids. Biophys J, 2006. 90(6): p. 1939–48.

64. Endres, D., et al., A reaction landscape identifies the intermediates critical for self-assembly of virus capsids and other polyhedral structures. Protein Science, 2005. 14: p. 1518–1525.

65. Wyman, J. and S.J. Gill, Binding and Linkage: Functional Chemistry of Biological Macromolecules. 1990, Herndon: University Science Books.

66. Schlicksup, C.J., et al., Hepatitis B virus core protein allosteric modulators can distort and disrupt intact capsids. eLife, 2018. 7: p. e31473.

67. Li, L., et al., Phase Diagrams Map the Properties of Antiviral Agents Directed against Hepatitis B Virus Core Assembly. Antimicrobial agents and chemotherapy, 2013. 57(3): p. 1505–8.

68. Zlotnick, A., et al., In vitro screening for molecules that affect virus capsid assembly (and other protein association reactions). Nature protocols, 2007. 2(3): p. 490–498.

69. Tang, G., et al., EMAN2: an extensible image processing suite for electron microscopy. J Struct Biol, 2007. 157(1): p. 38–46.

70. Scheres, S.H., RELION: implementation of a Bayesian approach to cryo-EM structure determination. J Struct Biol, 2012. 180(3): p. 519–30.

71. Zivanov, J., et al., New tools for automated high-resolution cryo-EM structure determination in RELION-3. Elife, 2018. 7: p. e42166.

72. Ilca, S.L., et al., Localized reconstruction of subunits from electron cryomicroscopy images of macromolecular complexes. Nature Communications, 2015. 6: p. 8843.

73. Huiskonen, J.T., Image processing for cryogenic transmission electron microscopy of symmetry-mismatched complexes. Bioscience reports, 2018. 38(2): p. BSR20170203.

74. Wynne, S.A., R.A. Crowther, and A.G. Leslie, The crystal structure of the human hepatitis B virus capsid. Mol Cell, 1999. 3(6): p. 771–80.

75. Adams, P.D., et al., PHENIX: a comprehensive Python-based system for macromolecular structure solution. Acta Crystallogr D Biol Crystallogr, 2010. 66(Pt 2): p. 213–21.

76. Emsley, P., et al., Features and development of Coot. Acta Crystallogr D Biol Crystallogr, 2010. 66(Pt 4): p. 486–501.

77. Moriarty, N.W., R.W. Grosse-Kunstleve, and P.D. Adams, electronic Ligand Builder and Optimization Workbench (eLBOW): a tool for ligand coordinate and restraint generation. Acta Crystallogr D Biol Crystallogr, 2009. 65(Pt 10): p. 1074–80.

78. Goddard, T.D., et al., UCSF ChimeraX: Meeting modern challenges in visualization and analysis. Protein Science, 2018. 27(1): p. 14–25.

79. Pettersen, E.F., et al., UCSF Chimera--a visualization system for exploratory research and analysis. J Comput Chem, 2004. 25: p. 1605–1612.

